# Automatic Inference of Sound Correspondence Patterns Across Multiple Languages

**DOI:** 10.1101/434621

**Authors:** Johann-Mattis List

## Abstract

Sound correspondence patterns play a crucial role for linguistic reconstruction. Linguists use them to prove language relationship, to reconstruct proto-forms, and for classical phylogenetic reconstruction based on shared innovations. Cognate words which fail to conform with expected patterns can further point to various kinds of exceptions in sound change, such as analogy or assimilation of frequent words. Here we present an automatic method for the inference of sound correspondence patterns across multiple languages based on a network approach. The core idea is to represent all columns in aligned cognate sets as nodes in a network with edges representing the degree of compatibility between the nodes. The task of inferring all compatible correspondence sets can then be handled as the well-known minimum clique cover problem in graph theory, which essentially seeks to split the graph into the smallest number of cliques in which each node is represented by exactly one clique. The resulting partitions represent all correspondence patterns which can be inferred for a given dataset. By excluding those patterns which occur in only a few cognate sets, the core of regularly recurring sound correspondences can be inferred. Based on this idea, the paper presents a method for automatic correspondence pattern recognition, which is implemented as part of a Python library which supplements the paper. To illustrate the usefulness of the method, we present how the inferred patterns can be used to predict words that have not been observed before.

## 1 Introduction

By comparing the languages of the world, we gain invaluable insights into human prehistory, predating the appearance of written records by thousands of years. The classical methods for historical language comparison, a collection of different techniques summarized under the term *comparative method* (Meillet, 1954; Weiss, 2015), date back to the early 19th century and have since then been constantly refined and improved (see Ross and Durie 1996 for details on the practical workflow). Thanks to the comparative method linguists have made groundbreaking insights into language change in general and into the history of many specific language families (Campbell and Poser, 2008) and external evidence has often confirmed the validity of the findings (McMahon and McMahon, 2005, 10-14). With increasing amounts of data, however, the methods, which are largely manually applied, reach their practical limits. As a result, scholars are now increasingly trying to automatize different aspects of the classical comparative methods (Kondrak, 2000; Prokić, Wieling, and Nerbonne, 2009; List, 2014).

One of the fundamental insights of early historical linguistic research was that – as a result of systemic changes in the sound system of languages – genetically related languages exhibit structural similarities in those parts of their lexicon which were commonly inherited from their ancestral languages. These similarities surface in form of *correspondence relations* between sounds from different languages in cognate words. English *th* [θ], for example, is usually reflected as *d* in German, as we can see from cognate pairs like English *think* vs. German *denken*, or English *thorn* and German *Dorn*. English *t*, on the other hand, is usually reflected as *z* [ts] in German, as we can see from pairs like English *toe* vs. German *Zeh*, or English *tooth* vs. German *Zahn*. The identification of these *regular sound correspondences* plays a crucial role in historical language comparison, serving not only as the basis for the proof of genetic relationship (Dybo and Starostin, 2008; Campbell and Poser, 2008) or the *reconstruction of proto-forms* (Hoenigswald 1960, 72-85, Anttila 1972, 229-263), but (indirectly) also for classical subgrouping based on shared innovations (which would not be possible without identified correspondence patterns).

With the beginning of this millennium, historical linguistics has witnessed an increased amount of attempts to quantify specific tasks of the traditional comparative method. Since then, scholars have repeatedly attempted to either directly infer regular sound correspondences across genetically related languages (Kondrak, 2009, 2003; Brown, Holman, and Wichmann, 2013; Kay, 1964) or integrated the inference into workflows for automatic cognate detection (Guy, 1994; List, 2012, 2014; List, Greenhill, and Gray, 2017). What is interesting in this context, however, is that almost all approaches dealing with regular sound correspondences, be it early formal – but classically grounded – accounts (Grimes and Agard, 1959; Hoenigswald, 1960) or computerbased methods (Kondrak, 2003, 2002; List, 2014) only consider sound correspondences between *pairs* of languages.

A rare exception can be found in the work of Anttila (1972, 229-263), who presents the search for regular sound correspondences across multiple languages as the basic technique underlying the comparative method for historical language comparison. Anttila’s description starts from a set of cognate word forms (or morphemes) across the languages under investigation. These words are then arranged in such a way that corresponding sounds in all words are placed into the same column of a matrix. The extraction of regularly recurring sound correspondences in the languages under investigation is then based on the identification of similar patterns recurring across different columns within the cognate sets. The procedure is illustrated in Figure 1, where four cognate sets in Sanskrit, Ancient Greek, Latin, and Gothic are shown, two taken from Anttila (1972, 246) and two added by me.

**Figure 1:**
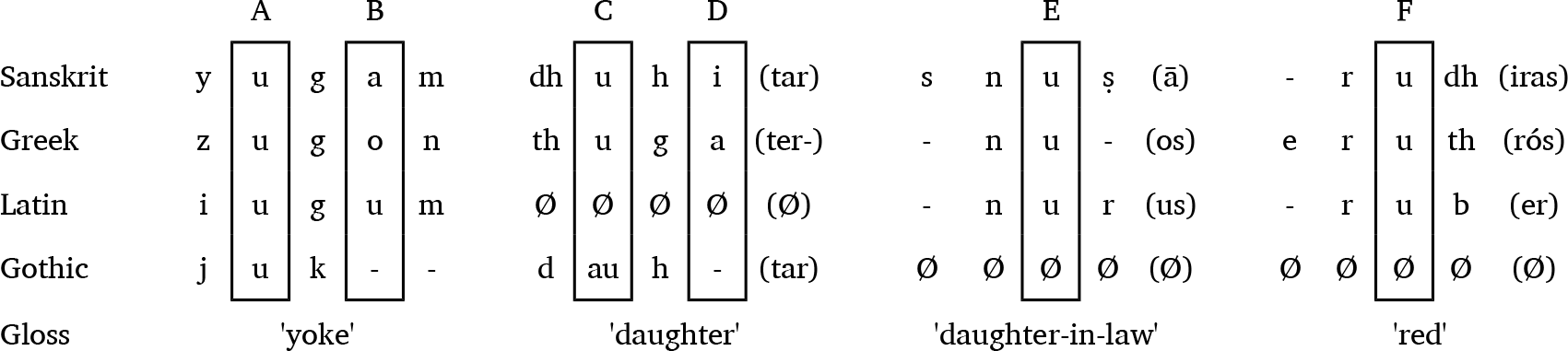
Regular sound correspondences across four Indo-European languages, illustrated with help of alignments along the lines of Anttila (1972: 246). In contrast to the original illustration, lost sounds are displayed with help of the dash “-” as a gap symbol, while missing words (where no reflex in Gothic or Latin could be found) are represented by the “Ø” symbol.

Two points are remarkable about Anttila’s approach. First, it builds heavily on the *phonetic alignment* of sound sequences,^1^, by which the sound sequences of words are arranged in a matrix in such a way that all corresponding sounds are placed in the same cell (List, 2014). Second, it reflects a concrete technique by which regular sound correspondences for multiple languages can be detected and employed as a starting point for linguistic reconstruction. If we look at the framed columns in the four examples in Figure 1, which are further labeled alphabetically, we can easily see that the patterns A, E, and F are remarkably similar. The only difference is that we miss data for Gothic in the patterns E and F, and, as a result, we don’t have *reflex sounds* (sounds in a given alignment column as reflected in a cognate word) for the full sound correspondence patterns in the respective columns. The same holds, however, for columns C, E, and F. Since A and C differ regarding the reflex sound of Gothic (*u* vs. *au*), they cannot be assigned to the same correspondence set at this stage, and if we want to solve the problem of finding the regular sound correspondences for the words in the figure, we need decide which columns in the alignments we assign to the same correspondence set, thereby ‘imputing’ missing sounds where we miss a reflex. Assuming that the “regular” pattern in our case is reflected by the group of C, E, and F, we can make *predictions* about the sounds missing in Gothic in E and F, concluding that, if ever we find the missing reflex in so far unrecognised sources of Gothic in the future, we would expect a *-au-* in the words for ‘daughter-in-law’ and ‘red’.^2^

We can easily see how patterns of sound correspondences across multiple languages can serve as the basis for multiple tasks in historical linguistics. First, we could use them to guess how a word that is missing in a given alignment would sound in that language, if it could be found. Since the task of identifying cognate words across multiple languages is very complex, and words may have drastically shifted their meanings, we could use the predictions to search for missing cognate forms in those areas of the lexicon which we have not considered before. Second, if two alignment columns are identical, they must reflect the same proto-sound, if alternative processes like borrowing can be excluded. Thus, similarly to the prediction of missing words in our cognate sets, we could use correspondence patterns to infer proto-forms, provided that parts of the data are already annotated.^3^ Third, we could use them to check linguistic claims about cognate words themselves: if it turns out that the aligned cognate sets proposed by linguists do not pattern into recurring correspondences across the languages under consideration, we can directly criticize both individual claims regarding word relations and general claims about the genetic relation of languages.

While it seems trivial to identify sound correspondences across multiple languages from the few examples provided in Figure 1, the problem can become quite complicated if we add more cognate sets and languages to the comparative sample. Especially the handling of *missing reflexes* for a given cognate set becomes a problem here, as missing data makes it difficult for linguists to decide which alignment columns to group with each other. This can already be seen from the examples given in Figure 1, where we have two possibilities to group the patterns A, C, E, and F.

The goal of this paper is to illustrate how a manual analysis in the spirit of Anttila can be automatized and fruitfully applied – not only in purely computational approaches to historical linguistics, but also in computer-assisted frameworks that help linguists to explore their data before they start carrying out painstaking qualitative comparisons (List, 2016). In order to illustrate how this problem can be solved computationally, the paper will first discuss some important general aspects of sound correspondences and sound correspondence patterns in Section 2, introducing specific terminology that will be needed in the remainder. In Section 3, we will see that the problem of finding sound correspondences across multiple languages can be modeled as the well-known *clique-cover problem* in an undirected network (Bhasker and Samad, 1991). While this problem is *hard* to solve in an exact way computationally,^4^ fast approximate solutions exist (Welsh and Powell, 1967) and can be easily applied. Based on these findings, the paper will introduce a fully automated method for the recognition of sound correspondence patterns across multiple languages (Section 4). This method is implemented in form of a Python library and can be readily applied to multilingual wordlist data as it is also required by software packages such as LingPy (List, Greenhill, and Forkel, 2017) or software tools such as EDICTOR (List, 2017). Section 5 will then illustrate how the method can be applied by testing how it performs in the task of predicting missing cognate words and missing proto-forms.

## 2 Preliminaries on Sound Correspondence Patterns

In the introduction, it was emphasized that the traditional comparative method is itself less concerned with regular sound correspondences attested for language pairs, but for all languages under consideration. In the following, this claim will be further substantiated, while at the same time introducing some major methodological considerations and ideas which are important for the development of the new method for sound correspondence pattern recognition.

### 2.1 From Sound Correspondences to Sound Correspondence Patterns

Sound correspondences are most easily defined for pairs of languages. Thus, it is straightforward to state that German [d] regularly corresponds to English [θ], that German [ts] regularly corresponds to English [t], and that German [t] corresponds to English [d]. We can likewise expand this view to multiple languages by adding another Germanic language, such as, for example, Dutch to our comparison, which has [d] in the case of German [d] and English [θ], [t] in the case of German [ts] and English [t], and [d] in the case of German [t] and English [d].

The more languages and examples we add to the sample, however, the more complex the picture becomes, and while we can state three (basic) patterns for the case of English, German, and Dutch, given in our example, we may get easily more patterns, due to secondary sound changes in the different languages, although we would still reconstruct only three sounds in the proto-language ([θ, t, d]). This is illustrated in Table 1, where Proto-Germanic forms containing *þ [θ], *t, and *d in different phonetic environments are contrasted with their descendant forms in German, English, and Dutch. The example shows that there is a one-to-*n* relationship between what we interpret as a proto-sound of the proto-language, and the regular correspondence patterns which we may find in our data. While we will reserve the term *sound correspondence* for pairwise language comparison, we will use the term *sound correspondence pattern* (or simply *correspondence pattern*) for the abstract notion of regular sound correspondences across a set of languages which we can find in the data.

**Table 1:**
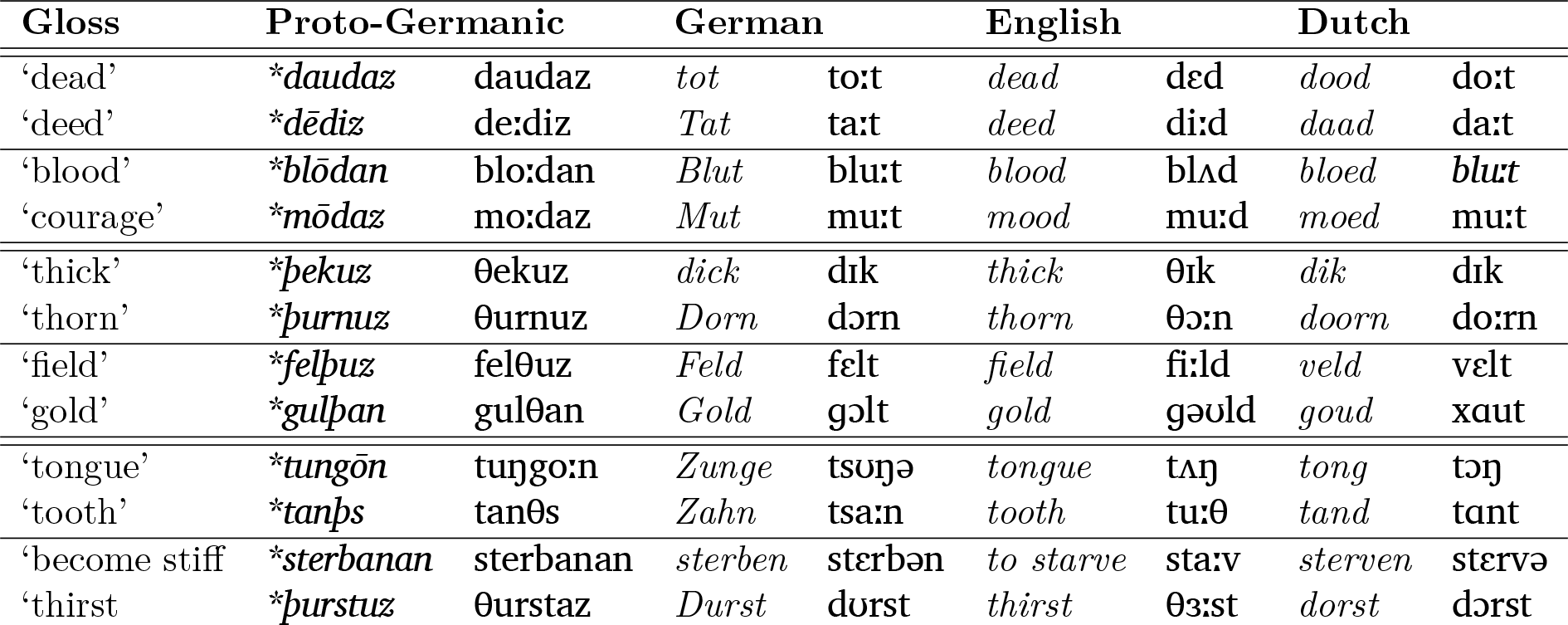
Comparing correspondence patterns for Proto-Germanic reflexes of *\*d-*, *\*þ-*, and *\*t-* in German, English, and Dutch (Germanic proto-forms follow Kroonen 2013).

### 2.2 Correspondence Patterns in the Classical Literature

Scholars like Meillet (1908, 23) have stated that the core of historical linguistics is not linguistic reconstruction, but the inference of correspondence patterns, emphasizing that ‘reconstructions are nothing else but the signs by which one points to the correspondences in short form’. ^5^ However, given the one-to-*n* relation between proto-sounds and correspondence patterns, it is clear, that this is not quite correct. Having inferred regular correspondence patterns in our data, our reconstructions will add a different level of analysis by further *clustering* these patterns into groups which we believe to reflect one single sound in the ancestral language.

That there are usually more than just one correspondence pattern for a reconstructed proto-sound is nothing new to most practitioners of linguistic reconstruction. Unfortunately, however, linguists do rarely list all possible correspondence patterns exhaustively when presenting their reconstructions, but instead select the most frequent ones, leaving the explanation of weird or unexpected patterns to comments written in prose. A first and important step of making a linguistic reconstruction system transparent, however, should start from an exhaustive listing of all correspondence patterns, including irregular patterns which occur very infrequently but would still be accepted by the scholars as reflecting true cognate words.

What scholars do instead is providing tables which summarise the correspondence patterns in a rough form, e.g., by showing the reflexes of a given proto-sound in the descendant languages in a table, where multiple reflexes for one and the same language are put in the same cell. An example, taken with modifications^6^ from Clackson (2007, 37), is given in Table 2. In this table, the major reflexes of Proto-Indo-European stops in 11 languages representing the oldest attestations and major branches of Indo-European, are listed. This table is a very typical example for the way in which scholars discuss, propose, and present correspondence patterns in linguistic reconstruction (Brown et al., 2011; Holton et al., 2012; Jacques, 2017; Beekes, 1995). The shortcomings of this representation become immediately transparent. Neither are we told about the frequency by which a given reflex is attested to occur in the descendant languages, nor are we told about the specific phonetic conditions which have been proposed to trigger the change where we have two reflexes for the same proto-sound. While scholars of Indo-European tend to know these conditions by heart, it is perfectly understandable why they would not list them. However, when presenting the results to outsiders to their field in this form, it makes it quite difficult for them to correctly evaluate the findings. A sound correspondence table may look impressive, but it is of no use to people who have not studied the data themselves.

**Table 2:**
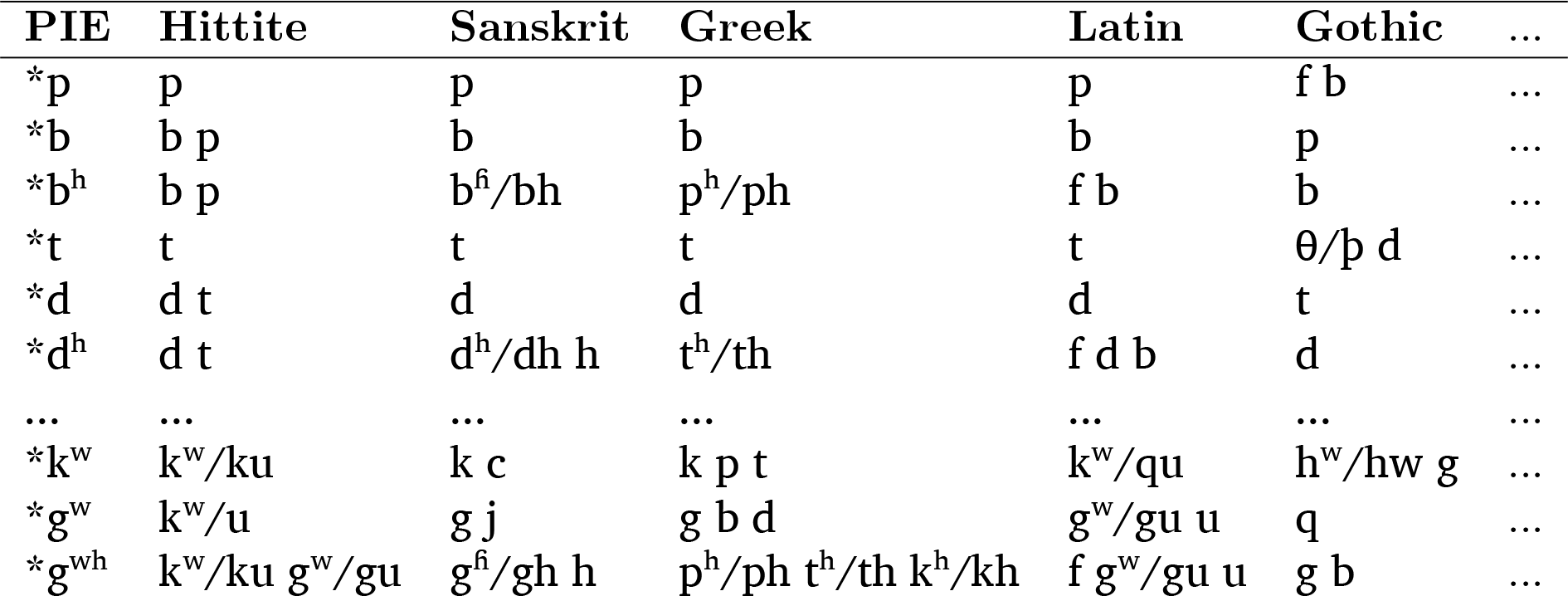
Sound correspondence patterns for Indo-European stops, following Clackson (2007, 37).

A further problem in the field of linguistic reconstruction is that scholars barely discuss workflows or procedures by which sound correspondence patterns can be *inferred*. For well-investigated language families like Indo-European or Austronesian, which have been thoroughly studied for more than hundred years (Blust, 1990), it is clear that there is no direct need to propose a heuristic procedure, given that the major patterns have been identified long ago and the research has reached a stage where scholarly discussions circle around individual etymologies or higher levels of linguistic reconstruction, like semantics, morphology and syntax.^7^ For languages whose history is less well known and where historical language reconstruction has not even reached a stage of reconstruction where a majority of scholars agrees, however, a procedure that helps to identify the major correspondence patterns underlying a given dataset, would surely be incredibly valuable.

### 2.3 Correspondence Patterns and Alignments

In order to infer correspondence patterns, the data must be available in *aligned* form (see Section 1), that is, we must know which of the sound segments that we compare across cognate sets are assumed to go back to the same ancestral segment. This is illustrated in Figure 2 where the cognate sets from Table 1 are presented in *aligned form*, with zero-matches (*gaps*) being represented as a dash ("-"), and with brackets indicating *unalignable parts* in the sequences, i.e., parts that cannot be aligned, since the differences are not due to regular sound change.^8^ Even if alignments are never mentioned in the entire book of Clackson (2007), the correspondence patterns shown in Table 2 directly reflect them, since each example that one could give for the data underlying a given correspondence pattern in the descendant languages would require the identification of unique sounds in each of the reflexes that confirm this pattern.

**Figure 2:**
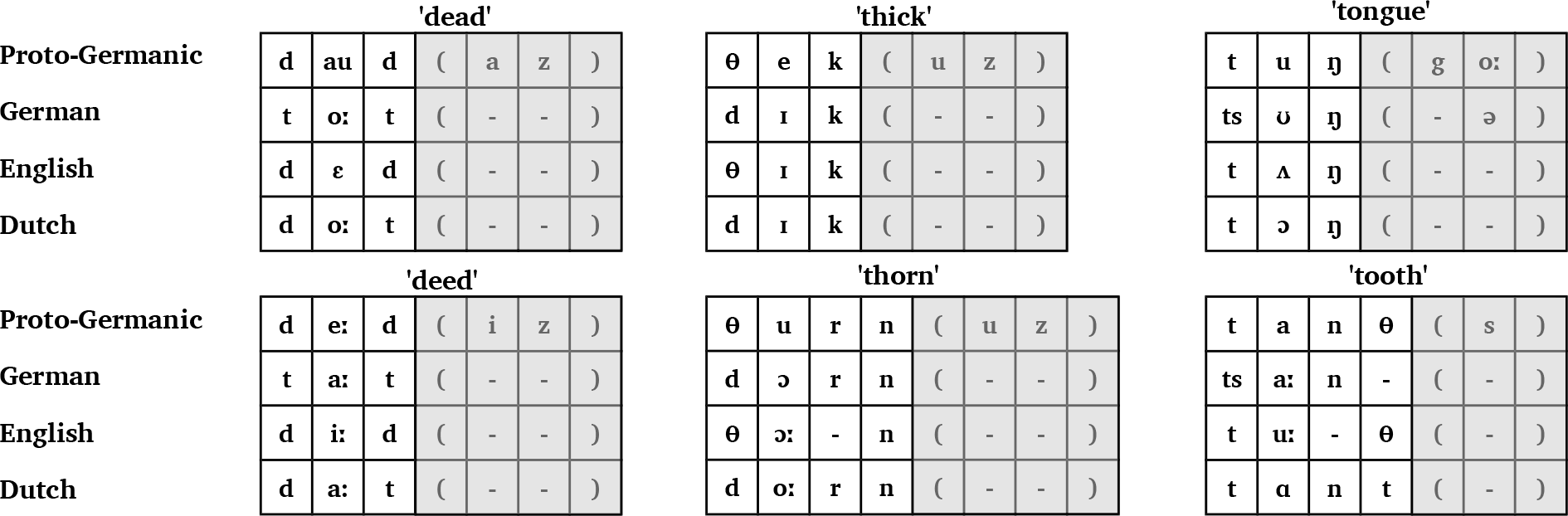
Alignment analyses of the six cognate sets from Table 1. Brackets around subsequences indicate that the alignments cannot be fully resolved due to secondary morphological changes.

Following evolutionary biology, a given column of an alignment is called an *alignment site* (or simply a *site*). An alignment site may reflect the same values as we find in a correspondence pattern, and correspondence patterns are usually derived from alignment sites, but in contrast to a correspondence pattern, an alignment site may reflect a correspondence pattern only incompletely, due to missing data in one or more of the languages under investigation. For example, when comparing German *Dorf* [dɔrf] ‘village’ with Dutch *dorp* [dɔrp], it is immediately clear that the initial sounds of both words represent the same correspondence pattern as we find for the cognate sets for ‘thick’ and ‘thorn’ given in Figure 2, although no reflex of their Proto-Germanic ancestor form *\*þurpa-* (originally meaning ‘crowd’, see Kroonen 2013, 553) has survived in Modern English.^9^ Thanks to the correspondence patterns in Table 1, however, we know that – if we project the word back to Proto-Germanic – we must reconstruct the initial with *\*þ-* ‘[θ], since the match of German *d-* and Dutch *d-* only occurs – if we ignore recent borrowings – only in correspondence patterns in which English has *th-*.

These “gaps” due to missing reflexes of a given cognate set are not the same as the gaps inside an alignment, since the latter are due to the (regular) loss or gain of a sound segment in a given alignment site, while gaps due to missing reflexes may either reflect processes of *lexical replacement* (List, 2014, 37f), or a preliminary stage of research resulting from insufficient data collections or insufficient search for potential reflexes. While we use the dash as a symbol for gaps in alignment sites, we will use the character Ø (denoting the empty set) to represent missing data in correspondence patterns and alignment sites. The relation between correspondence patterns in the sense developed here and alignment sites is illustrated in Figure 3, where the initial alignment sites of three alignments corresponding to Proto-Germanic *\*þ* [θ] are assembled to form one correspondence pattern.

**Figure 3:**
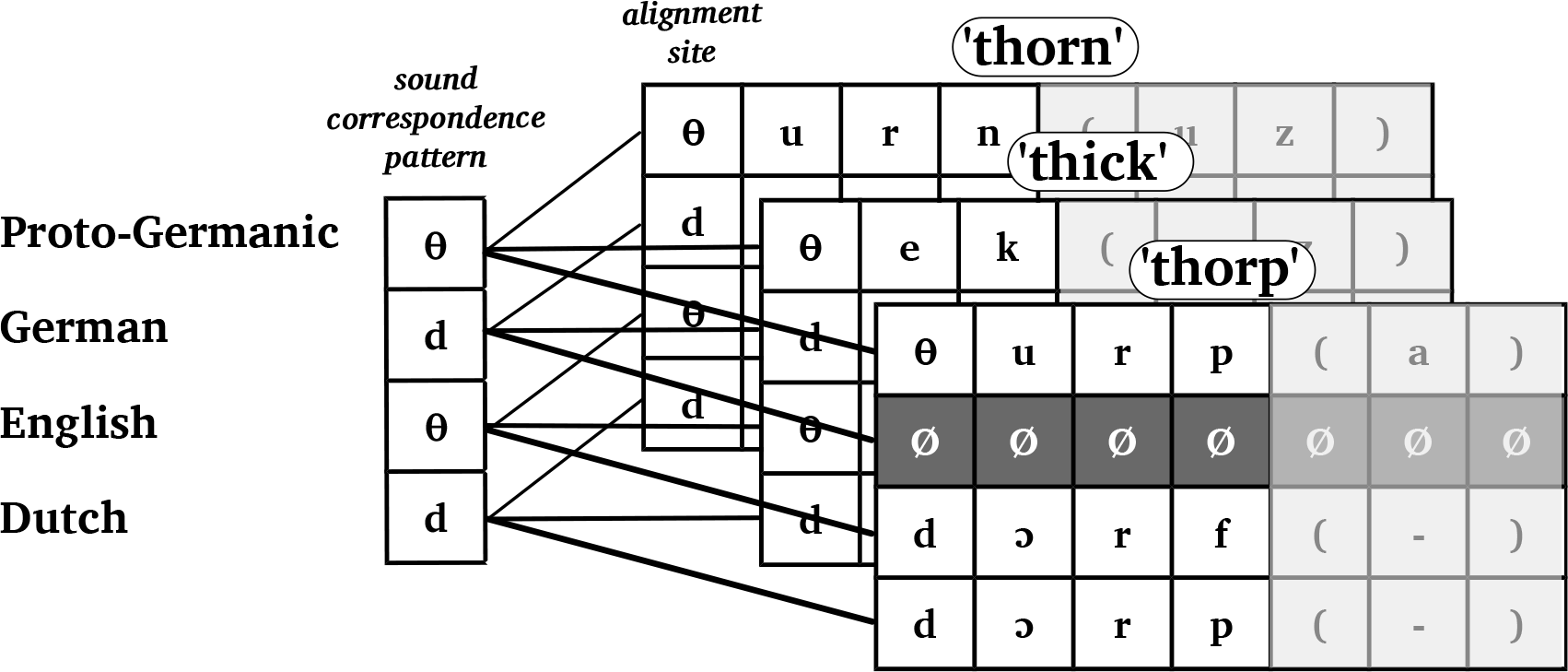
Alignment sites and correspondence patterns: While alignment sites are concrete representations of the presumed relations among cognate words, correspondence patterns are a further stage of abstraction.

## 3 Preliminary Thoughts on Correspondence Patterns Recognition

If we recall the problem we had in grouping the alignment sites E and F from Figure 1 with either A or C, we can see that the general problem of grouping alignment sites to correspondence patterns is their *compatibility*. If we had reflexes for all languages under investigation in all cognate sets, the compatibility would not be a problem, since we could simply group all identical sites with each other, and the task could be considered as solved. However, since it is rather an exception than the norm to have reflexes for all languages under consideration in a number of cognate sets, we will always find alternative possibilities to group our alignment sites in correspondence patterns. In the following, we will assume that two alignment sites are compatible, if they (a) share at least one sound which is not a gap symbol, and (b) do not have any conflicting sounds. This is illustrated in Figure 4 for our four alignment sites A, C, E, and F from Figure 1 above. As we can see from the figure, only two sites are incompatible, namely A and C, as they show different sounds for the reflexes in Gothic. Given that the reflex for Latin is missing in site C, we can further see that C shares only two sounds with E and F.

**Figure 4:**
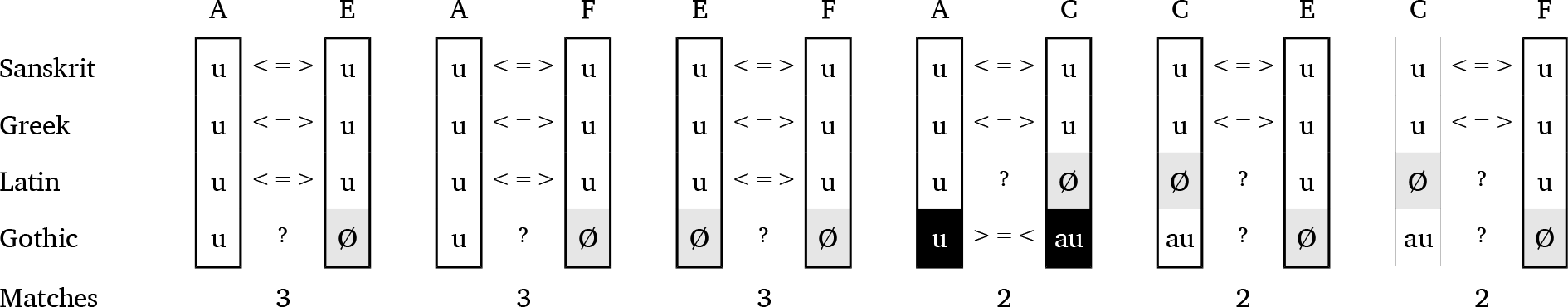
Assessing the compatibility of the four alignment sites from Figure 1.

Having established the notion of *alignment site compatibility*, it is straightforward to go a step further and model alignment sites in form of a *network*. Here, all sites in the data represent nodes (or vertices), and edges are only drawn between those nodes which are *compatible*, following the criterion of compatibility outlined in the previous section.^10^. Figure 5 illustrates how an alignment site network can be created from the compatibility comparison shown in Figure 4.

**Figure 5:**
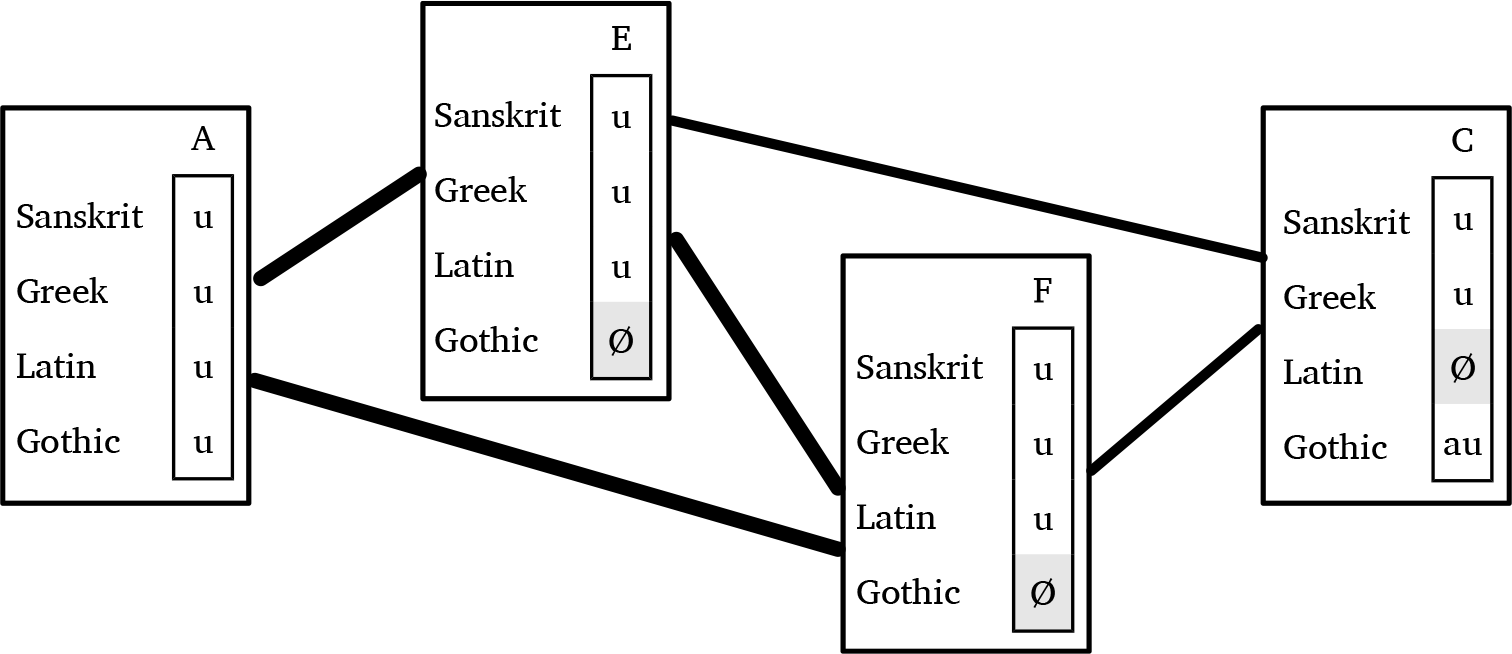
Representing alignment sites with help of a network. Edges are only drawn between compatible alignment sites. The width of the edges represents the number of matches per pair of alignment sites.

Having shown how the data can be modeled in form of a network, we can rephrase the task of identifying correspondence patterns as a *network partitioning task* with the goal to split the network into non-overlapping sets of nodes. Given that our main criterion for a valid correspondence pattern is full compatibility among all alignment sites of a given partition, we can further specify the task as a *clique partitioning task*. A *clique* in a network is ‘a maximal subset of the vertices [nodes] in an undirected network such that every member of the set is connected by an edge to every other’ (Newman, 2010, 193). Demanding that sound correspondence patterns should form a clique of compatible nodes in the network of alignment sites is directly reflecting the basic practice of historical language comparison as outlined by Anttila (1972), according to which a further grouping of incompatible alignment sites by proposing a proto-form would require us to identify a phonetic environment that could show incompatible sites to be complementary.

Parsimony dictates that – when partitioning our alignment site graph – we should try to minimize the number of cliques to which the different nodes are assigned. This is the *minimum clique cover problem* (Bhasker and Samad, 1991, 2). The minimum clique cover problem is a wellknown problem in graph theory and computer science, although it is usually more prominently discussed in form of its inverse problem^11^, the *graph coloring problem*. In the graph coloring problem, one tries to assign all those nodes in a graph to different clusters (i.e., to “color” them in different colors) which are directly connected (Hetland, 2010, 276). While the problem is generally known to be *NP-hard* (ibid.), fast approximate solutions like the Welsh-Powell algorithm (Welsh and Powell, 1967) are available. Using approximate solutions seems to be appropriate for the task of correspondence pattern recognition, given that we do not (yet) have formal linguistic criteria to favor one clique cover over another.^12^

## 4 An Automatic Method for Correspondence Pattern Recognition

The method for automatic correspondence pattern recognition requires that the data be coded for cognacy, and that all cognate sets be phonetically aligned. Thanks to recently proposed algorithms, these tasks can be carried out automatically,^13^ but to guarantee reliable results, it is useful to provide manually annotated data, or to manually correct data that was automatically analyzed in a first step.^14^

The general workflow of the method consists of three basic steps. In a first step, the alignments in the data are used to construct an *alignment site network* in which edges are drawn between compatible sites (A). The alignment sites are then partitioned into distinct non-overlapping subsets using an approximate algorithm for the minimum clique cover problem (B). In a final step, potential correspondence patterns are extracted from the non-overlapping subsets, and all individual alignment sites are re-assigned to those patterns with which they are compatible (C). The workflow is illustrated in Figure 6. In the following sections, I will provide more detailed explanations on the different stages.

**Figure 6:**
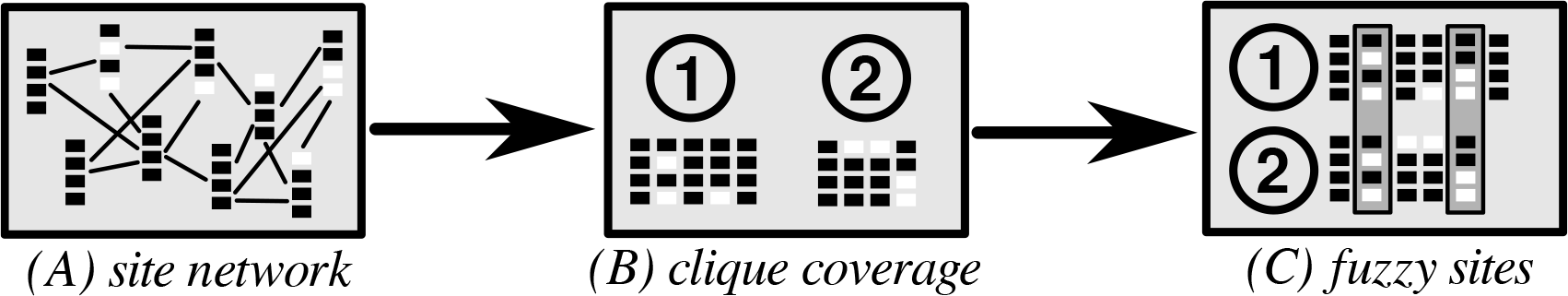
General workflow of the method for automatic correspondence pattern recognition.

### 4.1 Implementation, Input Format, and Output Format

The method has been implemented as a Python package that can be used as a plugin for the LingPy library for quantitative tasks in historical linguistics (List, Greenhill, and Forkel, 2017). The supplementary material offers precise instructions on how the software package can be installed and how the experiments can be replicated.

The input format for the method described here generally follows the input format employed by LingPy. In general, this format is a tab-separated text file with the first row being reserved for the header, and the first column being reserved for a unique numerical identifier. The header specifies the entry types in the data. Table 3 provides an example for the minimal data that needs to be provided to our method for automatic correspondence pattern recognition. In addition to the generally needed information on the identifier of each word (ID), on the language (DOCULECT), the concept or elicitation gloss (CONCEPT), the (not necessarily required) orthographic form (FORM), and the phonetic transcription provided in space-segmented form (TOKENS), the method requires information on the type of sound (consonant or vowel, STRUCTURE),^15^ the cognate set (COGID), and the alignment (ALIGNMENT).

**Table 3:**
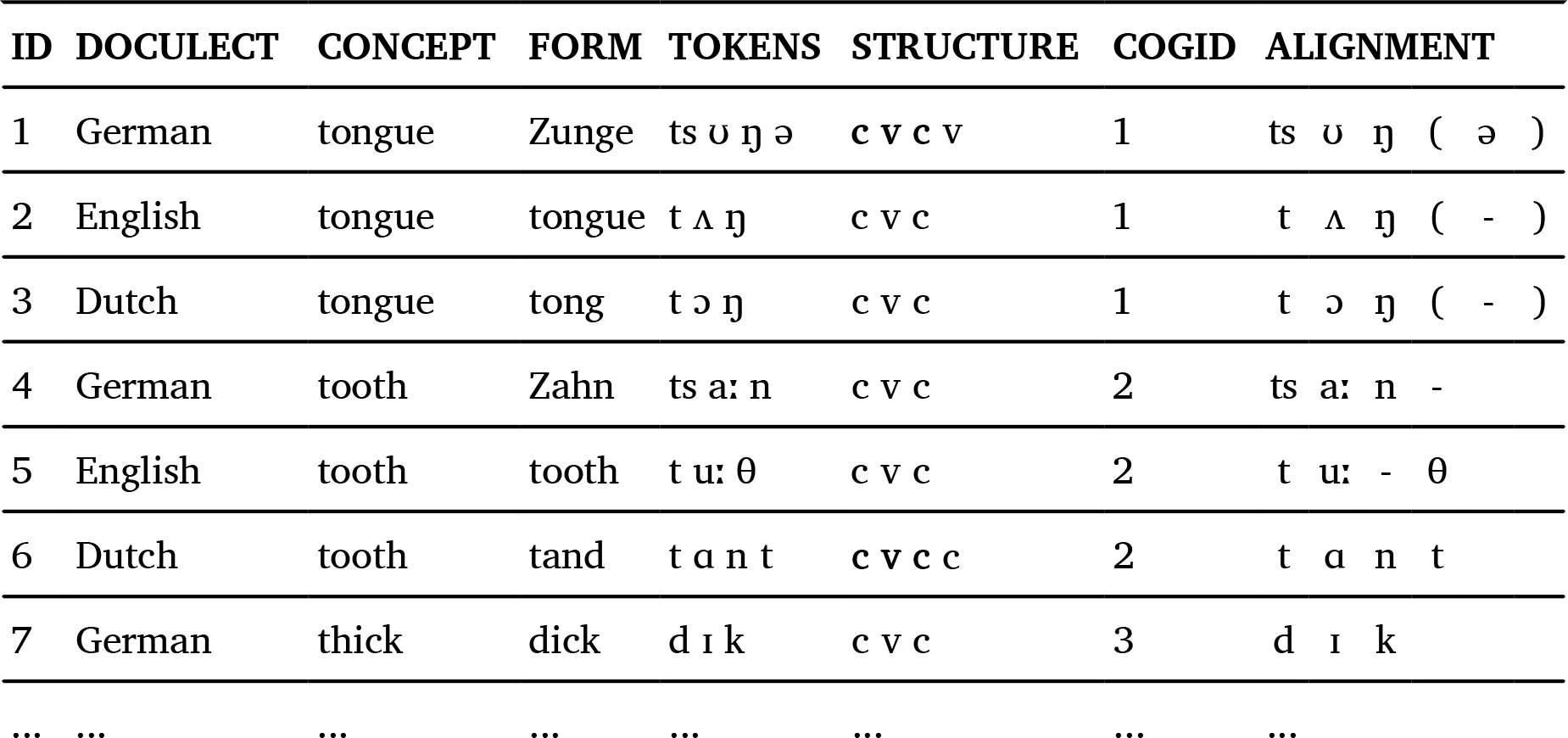
Input format with the basic values needed to apply the method for automatic correspondence pattern recognition.

The method offers different output formats, ranging from the LingPy wordlist format in which additional columns added to the original wordlist provide information on the inferred patterns, or in the form of tab-separated text files, in which the patterns are explicitly listed. The wordlist output can also be directly inspected in the EDICTOR tool, allowing for a convenient manual inspection of the inferred patterns.

### 4.2 Detailed Description of the Algorithm

As mentioned above, the method for correspondence pattern recognition consists of three stages. It starts with the reconstruction of an alignment site network in which each node represents a unique alignment site, and links between alignments sites are drawn if the sites are compatible, following the criterion for site compatibility outlined in Section 3 (A). It then uses a greedy algorithm to compute an approximate minimal clique cover of the network (B). All partitions proposed in stage (B) qualify as potentially valid correspondence patterns of our data. But the individual alignment sites in a given dataset may as well be compatible with more than one correspondence pattern.^16^ For this reason, the method iterates again over all alignment sites in the data and checks with which of the correspondence patterns inferred in stage (B) they are compatible. This procedure assigns each alignment site to at least one but potentially more different sound correspondence patterns (C).^17^

The clique cover algorithm (A) is an inverse version of the Welsh-Powell algorithm for graph coloring (Welsh and Powell, 1967). It starts by assigning all alignment sites in the data to distinct patterns, which are sorted in increasing order by the amount of missing data they contain. The algorithm then picks the first pattern and compares it with the set of all other patterns. If this first pattern is compatible with one of the other patterns, the two patterns will be merged into a new pattern that is then further compared with the remaining ones. After the iteration, the first pattern is added to the set of results, and the same procedure is repeated with the remaining patterns that have not yet been merged and remain in the queue until no patterns are left.

Since alignment sites may suffer from missing data, their assignment to particular correspondence patterns is not always unambiguous. The example alignment from Figure 1, for example, would yield two general correspondence patterns, namely u-u-u-au vs. u-u-u-u. While the assignment of the alignment sites A and C in the figure would be unambiguous, the sites E and F would be assigned to both patterns, since, judging from the data, we could not tell what correspondence pattern they represent in the end. In order to reflect the fuzziness of the pattern assignment, the method therefore requires an additional step. Here, we iterate again over all alignment sites and check with which of the inferred patterns they are compatible. The patterns, to which a given alignment site is assigned can further be ranked by counting the total amount of alignment sites with which they are compatible, thus allowing us to prefer only those site-to-pattern assignments that have a reasonable number of examples.

Figure 7 gives an artificial example that illustrates how the basic method infers the clique cover. Starting from the data in (A), the method assembles patterns A and B in (B) and computes their pattern, thereby retaining the non-missing data for each language in the pattern as the representative value. Having added C and D in this fashion in steps (C) and (D), the remaining three alignment sites, E-G are merged to form a new partition, accordingly, in steps (E) and (F). Step (G) reflects the re-assignment of individual alignment sites to the previously inferred patterns. In this example, all sites are only assigned to one pattern, but it is well possible, depending on the amount of missing data, that one site can be assigned to more than one pattern.

**Figure 7:**
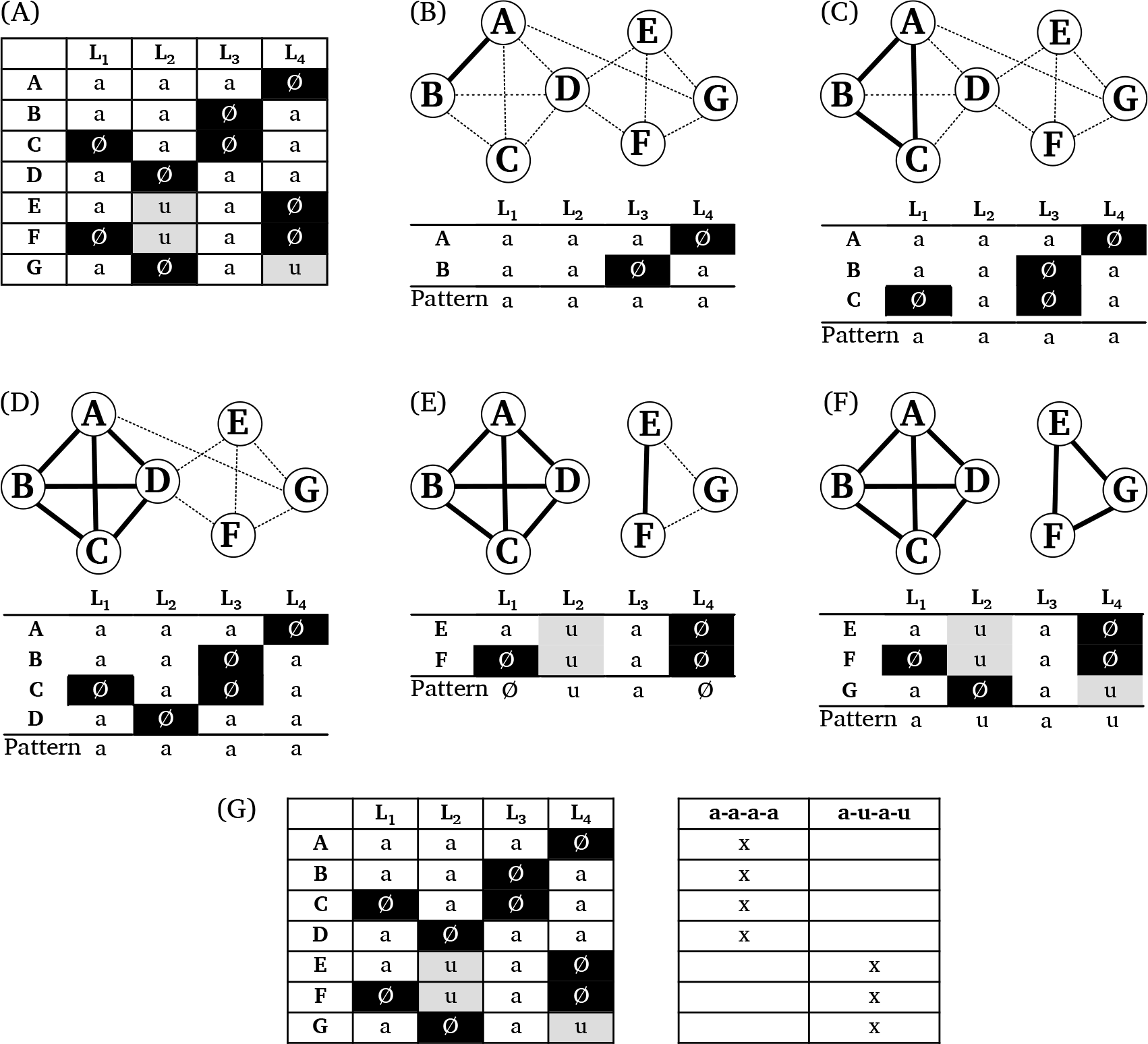
Example for the basic method to compute the clique cover of the data. (A) shows all alignment sites in the data. (B-D) show how the algorithm selects potential edges step by step in order to arrive at a first larger clique cover. (E-F) show how the second cover is inferred. In each step during which one new alignment site is added to a given pattern, the pattern is updated, filling empty spots. While there are two missing data points in (E), where only alignment sites E and F are merged, these are filled after adding G. (G) shows how patterns are re-assigned to individual alignment sites.

It is important to note that the originally selected pattern may change during the merge procedure, since missing spots can be filled by merging the pattern with a new alignment site (as also shown in Figure 7). For this reason, it is possible that this procedure, when only carried out one time, may not result in a true clique cover (in which all compatible alignment sites are merged). For this reason, at the end of the iteration, the algorithm checks if patterns exist that could be further combined, and repeats the procedure with the existing patterns until the resulting partitioning represents a true clique cover.

Pseudo code is in Algorithm 1 for the core function of the method for correspondence pattern detection. In a worst-case scenario in which all alignment sites will be assigned to distinct correspondence patterns, the algorithm requires 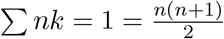 iterations in the while loop, where *k* represents the number of alignment sites in the data, so the general complexity of the algorithm is (𝒪*n*^2^). In applications to real-world-data, however, this worst-case scenario is never reached, and the method converges rather fast.

## 5 Testing the Method for Correspondence Pattern Recognition

The treatment of sound correspondence patterns presented in this study does not have predecessors in form of quantitative studies. As a result, no expert-annotated data listing all observable correspondence patterns for a certain language family exhaustively is available.^18^ and it is not possible to compare the suitability of this novel approach with expert-annotated gold standards, as it is usually done in similar studies in computational historical linguistics.

### Algorithm 1 Main part of the correspondence pattern detection method in pseudo-code.

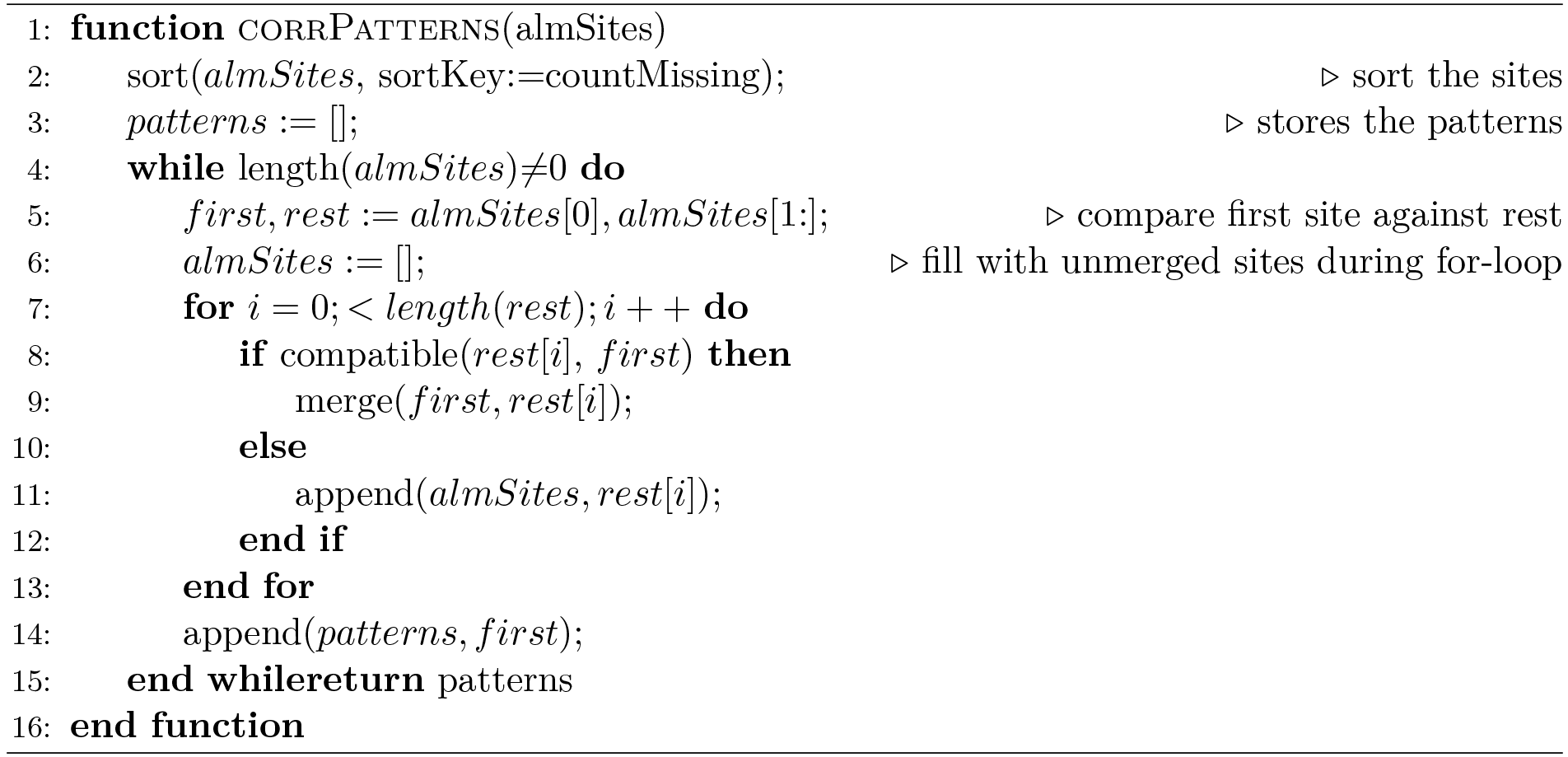

The lack of suitable gold standard data, however, does not mean that we cannot test the method for its suitability. Since the core service the method provides is to impute missing values in alignment sites resulting from cognate sets that are not reflected in all languages in a given dataset, we can easily design tests in which we test the power of the method to *predict* those missing values in controlled settings.

### 5.1 Data for Testing

Three different datasets were selected to test the method proposed in this study. The datasets were chosen with great care, since only a few of the many datasets offering manually coded cognate set also provide the cognate sets in aligned form. Apart from the data by Hill and List (2017) on Burmish languages (original data based on Huáng, Bùfán 1992), Walworth (forthcoming) on East Polynesian languages (original data based on Greenhill, Blust, and Gray 2008), and Hattori (1973) on Japanese languages (data in electronical form supplemented in List 2014), an additional dataset of 14 Chinese varieties originally published by Hóu (2004) was specifically modified and manually aligned for this study. While the two former datasets are classical wordlists that are further coded for cognacy and alignments,^19^ the Chinese data is based on a collection of 623 morphemes (reflected by a Chinese character each) whose pronunciation across the 14 dialects used in our sample was elicited by field workers. As a result, the amount of cognate sets with missing reflexes in this dataset is extremely low.

An overview of the datasets along with additional information regarding the data sources, the number of cognate sets, language varieties, and words in the data, is given in Table 4. Needless to say that all datasets are provided in the supplementary material accompanying this paper.

**Table 4:** Three test sets used in this study.

| Dataset | Source | Languages | Concepts | Words | Cognates |
| --- | --- | --- | --- | --- | --- |
| Burmish | Hill and List (2017) | 8 | 240 | 1819 | 798 |
| Chinese | Hóu (2004) | 14 | 623 | 8371 | 623 |
| Polynesian | Walworth forthcoming | 10 | 210 | 2427 | 1187 |
| Japanese | Hattori (1973) | 10 | 200 | 1986 | 454 |

### 5.2 General Characteristics

As a first illustrative test, the method was applied to the four datasets, and some basic statistics were calculated, these include the original number of alignment sites in the data, the number of patterns into which these sites were partitioned by the method, and the number of singleton patterns, i.e., patterns that are only reflected by one alignment site in the data. By dividing the number of alignment sites assigned to non-unique patterns by the number of all sites, we can further determine the proportion of “regular” correspondence patterns in a given dataset, assuming that a pattern is regular if it recurs in at least two different alignment sites.

The results of this analysis are summarized in Table 5. As we can see from this table, the number of correspondence patterns inferred by the algorithm is much lower than the number of alignment sites. This is, of course, not surprising, if we assume that the hypothesis that sound change is an overwhelmingly regular process, holds. However, across the datasets, we can find rather large differences with respect to the amount of singleton patterns, that is, patterns reflecting only one alignment site. That an alignment site is not compatible with any other site in the data can have different reasons. One reason are idiosyncratic sound changes, resulting, for example, from taboo, or from the assimilation of frequently used words. Another reason are errors in the data, resulting from incorrectly assigned cognates, alignments, or undetected borrowings. It is also possible that the data sample is too small, and that additional samples could be found, but have not been included in the data.

**Table 5:** Basic statistics after applying the correspondence pattern recognition method to the four datasets.

| Dataset | Align. Sites | Corr. Patterns | Singletons | Reg. Patterns |
| --- | --- | --- | --- | --- |
| Burmish | 1833 | 432 | 173 | 0.91 |
| Chinese | 2891 | 1341 | 966 | 0.67 |
| Polynesian | 1863 | 243 | 64 | 0.97 |
| Japanese | 1590 | 556 | 311 | 0.80 |

When comparing the proportion of “regular” patterns that are reflected by at least two alignment sites in the data across the datasets, the Chinese data shows the lowest proportion, with only 67% of all alignment sites being assigned to patterns that recur in the data. Given the intertwined history of the Chinese dialects, in which language contact among the dialect varieties played an important role, it is not necessarily surprising that the data looks less regular in general. As a manual inspection of the inferences reveals, the majority of the singleton alignment sites in the Chinese data could be assigned to one of the regular patterns if one of the reflexes would be ignored. Examples for these patterns are given in Table 6. On the other hand, we find some patterns which are largely irregular for specific reasons like taboo. An example is given in the same table with Chinese *niǎo* “bird” (pattern 679), which is reflected by nasal and dental initials across the Chinese dialects. As we know from older readings, the original reading had the initial [t], but it was later replaced by a nasal in some Chinese varieties to avoid homophony with the word for “penis”, which was most likely a metaphorically shifted from “bird”.

**Table 6:** Examples for idiosyncratic correspondence patterns in the Chinese dialects reflecting the major groups (Běijīng, Sūzhōu, Chángshā, Nánchāng, Méixiàn, Táoyuán, Guǎngzhōu, Fúzhōu, Táiběi).

| # | Cognates | BJ | SZ | CS | NC | MX | TY | GZ | FZ | TB |
| --- | --- | --- | --- | --- | --- | --- | --- | --- | --- | --- |
| 73 | 5 | t | d | t | t<sup>h</sup> | t<sup>h</sup> | t<sup>h</sup> | t | t | t |
| 484 | 1 | t | d | t | l | t<sup>h</sup> | t<sup>h</sup> | t | t | t |
| 654 | 1 | t | d | t | t<sup>h</sup> | t | t<sup>h</sup> | t | t | t |
| 239 | 6 | n | ɲ | ɲ | ɲ | ŋ | ŋ | n | n | l |
| 679 | 1 | n | ɲ | ɲ | ɲ | t | ŋ | n | ts | ts |
| 699 | 1 | n | ɲ | ɲ | ɲ | ŋ | ŋ | ŋ | g | g |

What we can see from the individual analyses of the different datasets is that the overall regularity of correspondence patterns does not necessarily reflect the time depth of the languages in a given dataset. Instead, correspondence patterns reflect different aspects of the data which have so far not been thoroughly investigated by researchers. The overwhelming regularity of the patterns in the Polynesian dataset, for example, is probably also due to the fact that the languages contain very small phoneme inventories, with no more than 17 different sounds on average (compared to Chinese dialects with about 35 sounds), while many idiosyncratic patterns in the Japanese data result from morphological differences which are difficult to handle in phonetic alignments.

### 5.3 Tests on Word Prediction

As mentioned briefly already in Section 1, correspondence patterns – once readily inferred – can provide hints regarding the potential pronunciation of missing cognates in an alignment. Since the method for correspondence pattern recognition imputes missing data in its core, it can also be used to predict how a given word should look in a given language if the reflex of the corresponding cognate set is missing. An example for the prediction of forms has been given above for the cognate set Dutch *dorp* and German *Dorf*. Since we know from Table 1 that the correspondence pattern of *d* in Dutch and German usually points to Proto-Germanic *\*þ*, we can propose that the English reflex (which is missing in Modern English, apart from place names) would start with *th*, if it was still preserved.^20^ Since the method for correspondence pattern recognition assigns one or more correspondence patterns to each alignment site, even if the site has missing data for a certain number of languages, all that needs to be done in order to predict a missing entry is to look up the alignment pattern and check the value that is proposed for the given language variety.

The test on word prediction was designed as follows: from each of the datasets, a certain number of cognate sets was randomly deleted, and the resulting data was then analyzed with help of the correspondence pattern recognition algorithm. In a second step, these inferred patterns were used to predict the cognate words which were deleted before. For the prediction, only the largest correspondence pattern was considered for the imputation, in order to avoid that multiple proposals for one sound could be made by the algorithm. For each dataset, three different proportions of words to be deleted were tested (25%, 50%, and 75%).^21^ For each proportion and dataset 1000 trials were tested and the results were averaged. To assess the accuracy of a predicted word, the proportion of correctly predicted sounds in the given word was estimated and divided by the total length of the word.

The results of this experiment are given in Table 7. In general, we can note that the prediction experiment works very well across all datasets for wordlists reduced by 25% and 50% of their words appearing in cognate sets, while the accuracy of prediction drastically drops in all datasets when removing up to 75% of the data. The only exception is the Polynesian dataset, where the difference in accuracy across the three experiments is only small, with a rather large standard deviation.

**Table 7:** Results of the test on word prediction, based on 1000 random samples for each subset of the data. The column Proportion reflects the different proportions of the data that was deleted during the experiments. Patterns refers to the average number of correspondence patterns inferred in each trial, and Reg. Patterns points to the proportion of alignment sites covered by patterns recurring at least twice.

| **Dataset** | **Proportion** | **Patterns** | **Reg. Patterns** | **Accuracy** |
| --- | --- | --- | --- | --- |
| Burmish | 25% | $231.68 \pm 6.86$ | $0.94 \pm 0.01$ | $0.59 \pm 0.02$ |
| | 50% | $165.55 \pm 6.08$ | $0.94 \pm 0.01$ | $0.53 \pm 0.02$ |
| | 75% | $99.33 \pm 5.71$ | $0.89 \pm 0.02$ | $0.37 \pm 0.03$ |
| Chinese | 25% | $1040.62 \pm 11.88$ | $0.81 \pm 0.01$ | $0.69 \pm 0.01$ |
| | 50% | $672.35 \pm 10.53$ | $0.95 \pm 0.00$ | $0.70 \pm 0.01$ |
| | 75% | $373.23 \pm 7.67$ | $0.97 \pm 0.00$ | $0.64 \pm 0.01$ |
| Japanese | 25% | $399.82 \pm 10.04$ | $0.89 \pm 0.01$ | $0.64 \pm 0.01$ |
| | 50% | $259.71 \pm 9.39$ | $0.93 \pm 0.01$ | $0.62 \pm 0.01$ |
| | 75% | $142.65 \pm 7.35$ | $0.92 \pm 0.01$ | $0.52 \pm 0.02$ |
| Polynesian | 25% | $127.30 \pm 5.38$ | $0.97 \pm 0.00$ | $0.81 \pm 0.01$ |
| | 50% | $89.10 \pm 5.51$ | $0.97 \pm 0.01$ | $0.82 \pm 0.01$ |
| | 75% | $51.37 \pm 4.69$ | $0.95 \pm 0.01$ | $0.80 \pm 0.03$ |

What may come as a surprise is that the reduction of the data by 25% and 50% does not seem to influence the accuracy of prediction in all datasets. On the contrary, in the Chinese and the Polynesian datasets, we find even slightly higher accuracy scores for the larger data reduction. At least in the Chinese data, the reason for this can be found in the large number of singleton patterns that deviate only in one reflex from regularly recurring patterns. If the data is reduced by 50%, the number of idiosyncratic patterns also drops, as we can see from the proportion of regularly recurring patterns given in the table. While these cover 95% of all alignment sites in the dataset reduced by 50%, their proportion drops to 81% when being reduced by only 25%, and is (as we have seen in Table 5) even lower when analyzing the whole dataset. If enough words are deleted from singleton patterns, like the ones shown in the examples in Table 6, the method for correspondence pattern recognition will assign them to the same clusters. As a result, the words whose pronunciation deviates will still be wrongly predicted, but the words that are not affected by individual sound changes will be predicted correctly, and since there are more regular words in the data, the overall prediction accuracy will increase.

When comparing the differences in the scores across the four datasets, we can also see that the overall “regularity” of the data, as measured by the amount of patterns that recur more than two times, is not a good predictor of the success of the prediction quality. The Burmish data, for example, has rather high rates of pattern regularity, but performs worse in prediction than the other datasets. It is clear that the number of singleton patterns that only reflect one alignment site in a dataset will have a direct impact on the word prediction quality, since only patterns that recur at least two times in the data can be used for prediction. But this is not the only factor influencing the prediction quality. Ambiguous alignment sites that can be assigned to more than one pattern may, for example, likewise produce erroneous predictions. For the time being, we cannot offer a full account on all different factors that might influence prediction quality. More studies on different datasets will be needed to increase our knowledge in the future.

The fact that the prediction accuracy does not seem to improve or may even drop when more data is retained in our experiments is important for further applications of the methods (for example when carrying out field work or when searching for missing cognate sets), as it shows that we can reduce the amount of time spent on manual annotation substantially when annotating datasets for historical linguistics. Linguists could, for example, annotate half of their data manually and then use our method to impute potentially missing cognates in their data. If actual words that were not annotated in the first run turn out to have the same form as words predicted by our algorithm, this would be a very strong argument that they are really cognate. Another example would be guided field work for the purpose of historical language comparison. If insufficient amounts of data have been collected, scholars can use the prediction method to predict the most likely forms for certain cognate sets and use them to ease the elicitation of the relevant forms when asking new informants.

### 5.4 Examples

Table 8 gives some examples illustrating the scoring procedure and typical failures of the method, again illustrated for the Chinese dataset.^22^ In cognate set 687, we find one correctly predicted form for Chángshā, and one incorrectly predicted tone for the Jínán form. As we can see from the frequencies of alignment sites supporting the proposed pattern, the inferred pattern clusters only two alignment sites. As a result, it is not surprising that a wrong tone is proposed. The wrong form for Měixiān in cognate set 319 is due to a wrong clustering of the cognate set with the irregular cognate set 654 listed above in Table 6. Since the Měixiān word was deleted in the experiment, the whole pattern is compatible with pattern # 73 in the table, which predicts that the Měixiān form should start with tʰ. In cognate set 518, we can see that the method fails to propose a valid sound for the form in Wēnzhōu for the second and the third site in the alignment, given that these sites are assigned to patterns in which no sound for Wēnzhōu could be imputed.

**Table 8:**
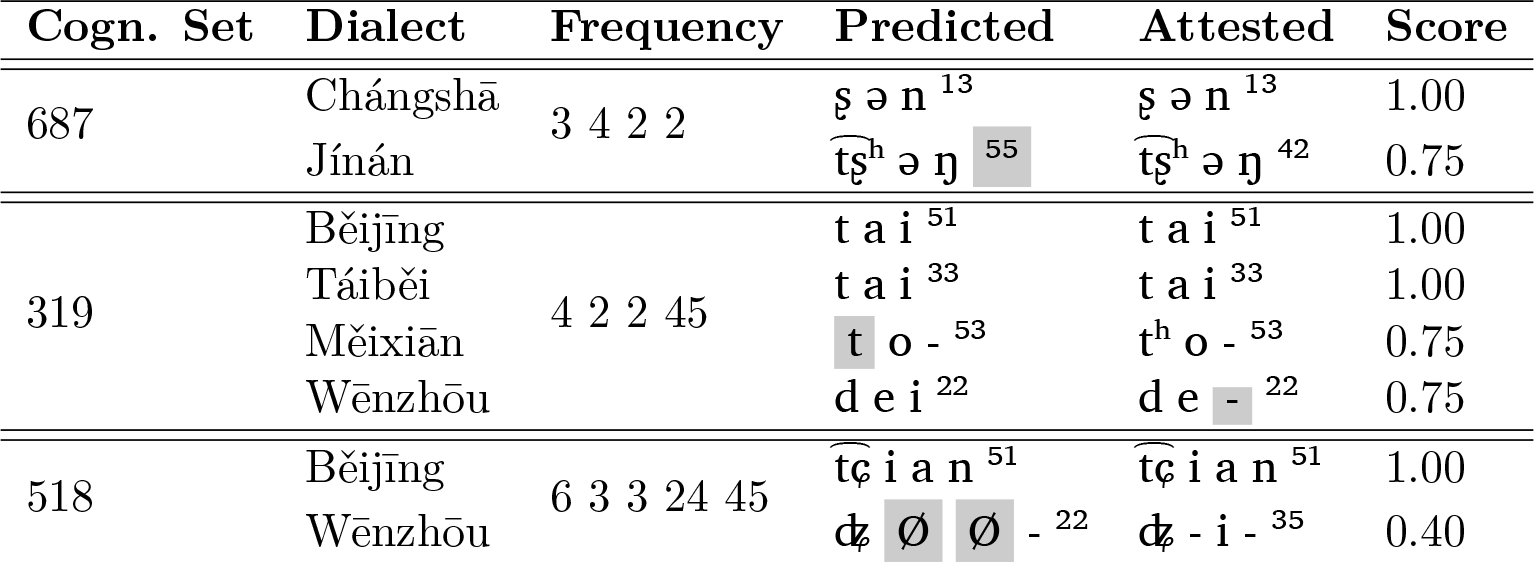
Examples for the word prediction experiment for the Chinese data. The column Frequency lists the size of the inferred patterns for each position of the predicted word form. The score is calculated by dividing the number of correctly predicted sounds by the total number of sounds.

While the success or failure of the prediction experiments can help us to improve the method in the future, we can also illustrate how the analysis can aid in practical work on linguistic reconstruction. This example will gain be based on the Chinese data, since it has the advantage of offering quick access to Middle-Chinese reconstructions. Since Middle Chinese is only partially reconstructed on the basis of historical language comparison, and mostly based on written sources, such as ancient rhyme books and rhyme tables (Baxter, 1992), the reconstructions are not entirely dependent on the modern dialect readings, which is a great advantage for testing the consequences of the correspondence pattern analysis.

In Table 9, patterns inferred by the method for correspondence pattern recognition for a reduced number of dialects (one of each major subgroup) have been listed. The examples can all be reconstructed to a dental stop in Middle Chinese (*t, *t^h^ or *d). If we only inspect reflexes of Middle Chinese *d in the data, we can see that the initial is reflected in seven different patterns in our data. Four of these patterns, however, occur only one time (# 719, # 1096, # 484, and # 654), and if we exclude the reflexes for Méixiàn (# 719, # 654), Táiběi (# 1096), and Nánchāng (# 484), respectively, we can assign # 719 and # 1096 to # 718 and # 484 and # 654 to # 73. In patterns # 718 and # 747, only Fúzhōu shows a different reflex. Since we have forms that are homophones in Middle Chinese in both correspondence patterns (糖 in # 747 and 堂 i # 718 were both pronounced as *dam in Middle Chinese), we cannot find a conditioning context that would explain this difference from the perspective of Middle Chinese alone. We know, however, that the Mǐn dialects (to which Fúzhōu belongs) reflect features that are more archaic than Middle Chinese. In this case, the difference between the patterns is regularly reflecting the difference between plain voiced and breathy voiced initials in the ancestor of the Mǐn dialects, with the latter going back to complex onsets in Old Chinese, the predecessor of all Chinese dialects (Baxter and Sagart, 2014, 171f). Furthermore, if we compare the patterns # 747 and # 73 directly, we can see that, although only Súzhōu has a direct reflex of the original voiced sound in Middle Chinese, we can still find its traces in the different correspondence patterns, since Běijīng and Guǎngzhōu have contrastive outcomes in both patterns ([tʰ] vs. [t]). When inspecting the tones which are reconstructed for the different words in Middle Chinese, we can easily find a conditioning context why the reflexes differ. The *píng* (flat) tone category in Middle Chinese correlates with aspiration, while the other tone categories correlate with devoicing in the three dialects.^23^ If we had no knowledge of Middle Chinese, it would be harder to understand that both patterns correspond to the same proto-sound, but once assembled in such a way, it would still be much easier for scholars to search for a conditioning context that allows them to assign the same proto-sound to the two patterns in questions.

**Table 9:** Contrasting inferred correspondence patterns with Middle Chinese reconstructions (MC) and tone patterns (MC Tones: P: píng (flat), S: shǎng (rising), Q: qù (falling), R: rù (stop coda)) for representative dialects of the major groups (Běijīng, Sūzhōu, Chángshā, Nánchāng, Méixiàn, Táoyuán, Guǎngzhōu, Fúzhōu, Táiběi).

| # | Cogn. | MC | MC Tones | BJ | SZ | CS | NC | MX | TY | GZ | FZ | TB |
| --- | --- | --- | --- | --- | --- | --- | --- | --- | --- | --- | --- | --- |
| 30 | 13 | *t | PSQR | t | t | t | t | t | t | t | t | t |
| 41 | 9 | *th | PSQR | t<sup>h</sup> | t<sup>h</sup> | t<sup>h</sup> | t<sup>h</sup> | t<sup>h</sup> | t<sup>h</sup> | t<sup>h</sup> | t<sup>h</sup> | t<sup>h</sup> |
| 747 | 3 | *d | P | t<sup>h</sup> | d | t | t<sup>h</sup> | t<sup>h</sup> | t<sup>h</sup> | t<sup>h</sup> | t<sup>h</sup> | t<sup>h</sup> |
| 718 | 2 | *d | P | t<sup>h</sup> | d | t | t<sup>h</sup> | t<sup>h</sup> | t<sup>h</sup> | t<sup>h</sup> | t | t<sup>h</sup> |
| 719 | 1 | *d | P | t<sup>h</sup> | d | t | t<sup>h</sup> | t | t<sup>h</sup> | t<sup>h</sup> | t | t<sup>h</sup> |
| 1096 | 1 | *d | P | t<sup>h</sup> | Ø | t | t<sup>h</sup> | t<sup>h</sup> | t<sup>h</sup> | t<sup>h</sup> | t | t |
| 73 | 5 | *d | QR | t | d | t | t<sup>h</sup> | t<sup>h</sup> | t<sup>h</sup> | t | t | t<sup>h</sup> |
| 484 | 1 | *d | R | t | d | t | l | t<sup>h</sup> | t<sup>h</sup> | t | t | t<sup>h</sup> |
| 654 | 1 | *d | S | t | d | t | t<sup>h</sup> | t | t<sup>h</sup> | t | t | t<sup>h</sup> |

The example shows that, as far as the Middle Chinese dental stops are concerned, we do not find explicit exceptions in our data, but can rather see that multiple correspondence patterns for the same proto-sound may easily evolve. We can also see that a careful alignment and cognate annotation is crucial for the success of the method, but even if the cognate judgments are fine, but the data are sparse, the method may propose erroneous groupings.

In contrast to manual work on linguistic reconstruction, where correspondence patterns are never regarded in the detail in which they are presented here, the method is a boost, especially in combination with tools for cognate annotation, like EDICTOR, to which we added a convenient way to inspect inferred correspondence patterns interactively (see the example in Appendix A). Since linguists can run the new method on their data and then directly inspect the consequences by browsing all correspondence patterns conveniently in the EDICTOR, the method makes it a lot easier for linguists to come up with first reconstructions or to identify problems in the data.

## 6 Conclusion and Outlook

This study has presented a new method for the inference of sound correspondence patterns in multi-lingual wordlists. Thanks to its integration with popular software packages, the method can be easily applied, both within automated, or computer-assisted workflows. The usefulness of the method was illustrated by showing how it can be used to predict missing words in linguistic datasets. The method, however, has a lot of additional potential. Since the method can impute words not attested in existing languages, it could likewise be used for the automatic reconstruction of proto-forms, the identification of cognates, or the assessment of the general regularity of a given dataset. The value of the method is not that it reveals new things about a given language, but rather that it helps to assess how well a given dataset has been analyzed before. By helping to improve the quality of existing and future datasets in historical linguistics, we further hope that the method will on the long run also contribute to new and important findings about the past of our world’s languages.

## Supplementary Material

The supplementary material accompanying this paper contains the code and all instructions needed to repeat the experiments described in this paper. The original package for correspondence pattern detection is publicly available from GitHub under https://github.com/lingpy/lingrex. The package providing the supplementary material with results and instructions for running the code is also available via GitHub under https://github.com/lingpy/ correspondence-pattern-paper.

## Appendix A: Inspecting Correspondence Patterns in EDICTOR

The following screenshots shows how the modified version of the EDICTOR allows for an enhanced inspection of sound correspondence patterns inferred by the method.

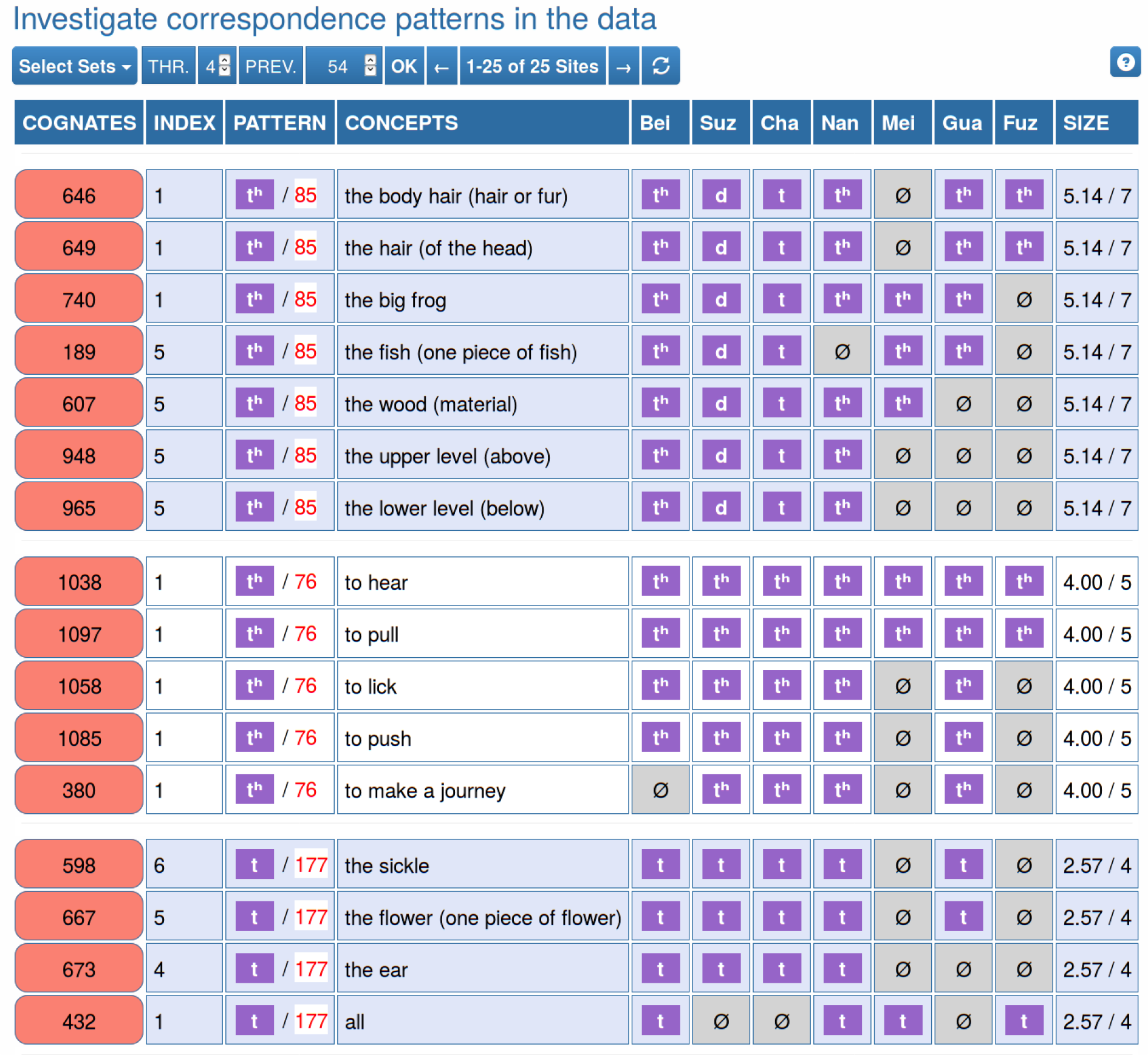

## Footnotes

1 This concept was only recently adapted in linguistics (Covington, 1996; Kondrak, 2000; List, 2014), building heavily on approaches in bioinformatics and computer science (Wagner and Fischer, 1974; Needleman and Wunsch, 1970), although it was implicitly always an integral part of the methodology of historical language comparison (compare Fox 1995, 67f, Dixon and Kroeber 1919)

2 As pointed out by the anonymous reviewer, Gothic *ráuþs* is indeed a reflex of ‘red’ (Wright, 1910, 340).

3 But even if correspondence patterns are not identical, they could be assigned to the same proto-sound, provided that one can show that the differences are conditioned by phonetic context. This is the case for Gothic *au* [o] in pattern *C*, which has been shown to go back to *u* when preceding *h* (Meier-Brügger, 2002, 210f). As a result, scholars usually reconstruct Proto-Indo-European *\*u* for A, C, E, and F.

4 Both the clique-cover problem and its inverse problem, the graph coloring problem, have been shown to be *np-complete* (Bhasker and Samad, 1991).

5 My translation, original text: ‘Les «restitutions» ne sont rien autre chose que les signes par lesquels on exprime en abrégé les correspondances’.

6 We added phonetic transcriptions, preceding the original sound given by the author, separated by a slash.

7 For examples, compare the very detailed etymological discussions by Meier-Brügger (2002, 173-187).

8 Scholars at times object to this claim, but it should be evident, also from reading the account by Anttila (1972) mentioned above, that without alignment analyses, albeit implicit ones that are never provided in concrete, no correspondence patterns could be proposed.

9 Old English still has the word *þorp*, but in Modern English, we only find *thorp* it in names.

10 We can further weight the edges in the alignment site network, for example, by using the number of matching sounds (where no missing data is encountered) to represent the strength of the connection (but we will disregard weighting in the approach presented here)

11 The inverse problem of a given problem in graph theory provides a solution to the original problem for a graph in which the original edges are deleted and nodes formerly unconnected are connected.

12 We should furthermore bear in mind that an optimal resolution of sound correspondence patterns for linguistic purposes would additionally allow for uncertainty when it comes to assigning a given alignment site to a given sound correspondence pattern. If we decided, for example, that the pattern C in Figure 5 could by no means cluster with E and F, this may well be premature before we have figured out whether the two patterns (u-u-u-u vs. u-u-u-au) are *complementary* and what phonetic environments explain their complementarity.

13 For automatic cognate detection, compare for example List (2014), List, Greenhill, and Gray (2017), Arnaud, Beck, and Kondrak (2017), and Jäger, List, and Sofroniev (2017), and for automatic phonetic alignment, compare Prokić, Wieling, and Nerbonne (2009) and List (2014).

14 For manual annotation of cognates and alignments, compare List (2017).

15 The values passed to the STRUCTURE column can be arbitrarily filled. When running the analysis, they are used to identify those positions in the alignments which should be analysed separately, i.e., they will be considered as a useful pre-partitioning of the alignment sites.

16 Compare, e.g., site E in Figure 1, which is both compatible with the pattern *u-u-u-u* reflected by the site A,

17 By further weighting and sorting the fuzzy patterns to which a given site has been assigned, the number of fuzzy alignment sites can be further reduced.

18 As we have seen in Figure 2, scholars list major sound correspondences across multiple languages, but they do not show individual patterns for aligned cognate sets.

19 All datasets are coded for partial cognates and across semantic categories.

20 We ignore deliberately in this context that the alternative of the correspondence in Dutch and German would be a borrowing from Dutch, Frisian, or English to German.

21 Depending on the specific distribution of cognates in the individual data, these proportions could vary in each run.

22 This dataset and the detailed predictions are available from the supplementary material as files `<chinese25.tsv>` (wordlist) and `predictions-chinese25.txt`.

23 This phenomenon most likely goes back to an earlier phonation contrast between the first (*píng*) tone in Middle Chinese and the other tones.

